# MALVA: genotyping by Mapping-free ALlele detection of known VAriants

**DOI:** 10.1101/575126

**Authors:** Giulia Bernardini, Paola Bonizzoni, Luca Denti, Marco Previtali, Alexander Schönhuth

## Abstract

The amount of genetic variation discovered and characterized in human populations is huge, and is growing rapidly with the widespread availability of modern sequencing technologies. Such a great deal of variation data, that accounts for human diversity, leads to various challenging computational tasks, including variant calling and genotyping of newly sequenced individuals. The standard pipelines for addressing these problems include read mapping, which is a computationally expensive procedure. A few mapping-free tools were proposed in recent years to speed up the genotyping process. While such tools have highly efficient run-times, they focus on isolated, bi-allelic SNPs, providing limited support for multi-allelic SNPs, indels, and genomic regions with high variant density.

To address these issues, we introduce `MALVA`, a fast and lightweight mapping-free method to genotype an individual directly from a sample of reads. `MALVA` is the first mapping-free tool that is able to genotype multi-allelic SNPs and indels, even in high density genomic regions, and to effectively handle a huge number of variants such as those provided by the 1000 Genome Project. An experimental evaluation on whole-genome data shows that `MALVA` requires one order of magnitude less time to genotype a donor than alignment-based pipelines, providing similar accuracy. Remarkably, on indels, `MALVA` provides even better results than the most widely adopted variant discovery tools.

## 1 Introduction and Related Work

The discovery and characterization of sequence variations in human populations is crucial in genetic studies. A prime challenge is to efficiently analyze the variations of a freshly-sequenced individual with respect to a reference genome and the available genomic variations data. To reach this goal, the standard pipeline includes aligning sequenced reads with softwares like `BWA` [19] and `Bowtie` [17] and then calling the genotypes (*e.g*., with `GATK` [22] or `bcftools` [20]); such an approach, though, can be highly time consuming, thus impractical for clinical applications, where time is often an issue. Typically in diploid organisms variant calling requires SNPs and indel detection and the identification of the pairs of alleles for each position of the studied genome, called genotype.

Recent tools for genotyping and variant calling like `Graphtyper` [11] and `vg` [31], which are based on a graph representation of a pan-genome to avoid biases introduced by considering only the information included in a set of genomes [5], are nevertheless heavy in both computational space and time. Moreover, the size of indexes of variation graphs may be subjected to an exponential growth in the number of variants included, and indexes typically require a great deal of computational resources to be updated with newly discovered variants. When the task is to call the genotype in positions where variants have been previously annotated, alignment-free methods come to the aid. Recent mapping-free genotyping tools such as `LAVA` [30] and `VarGeno` [34] are word-based methods that, given a list of known SNP loci, call SNPs as either mutant or wild-type up to an order of magnitude faster than the usual alignment-based methods. A major shortcoming of these tools is the large memory requirement, that can easily exceed hundreds of GB of RAM. Their strategy is to create a dictionary for both the reference genome and the SNP list that maps each *k*-mer to the positions at which it appears, and then to call variants from the reads by evaluating *k*-mers frequency. `FastGT` [27] is yet another *k*-mer-based method to genotype sequencing data: it strongly relies on a pre-compiled database of bi-allelic SNVs and corresponding *k*-mers, obtained by subjecting the *k*-mers that overlap known SNVs to several filtering steps. Such filters remove from the database the SNPs for which unique *k*-mers (*i.e*., not occurring elsewhere in the reference genome) are not observed, those that are closely located (*i.e*., that are less than k bases apart), and others: after the filtering steps, only 64% of bi-allelic SNVs survive and are therefore identifiable. These tools implement strategies to represent and analyze SNPs that improve the time performance but, on the other hand, do not allow to model indels and close variants.

Short insertions and deletions of nucleotides (*indels*) are believed to represent around 16% to 25% of human genetic polymorphism [24]. The presence of indels can be associated with a number of human diseases [26, 13, 16]: for instance, cystic fibrosis [26], lung cancer [29], Mendelian disorders [21] and Bloom syndrome [14] are all known to be closely correlated to indels. Indels are particularly challenging to call from NGS data, because mapping is more difficult when the reads overlap with indels [25].

In this paper we introduce `MALVA`, a rapid, lightweight, alignment-free method to genotype known (*i.e*., previously characterized) variants, including indels and close SNPs, in a sample of reads. `MALVA` is a word-based method: each allele of each known variant is assigned a *signature* in the form of a set of *k*-mers, which allows to efficiently model indels and close variants. The genotypes will be called according to the frequency of such signatures in the input reads. Based on the well-known Bayes’ formula, we also design a new rule to genotype multi-allelic variants (*i.e*., variants such that more than one alternate allele is known): even if such variants are trickier to genotype than bi-allelic ones, we are still able to achieve high precision and recall, as revealed in the real-data experiments we conducted. `MALVA` directly analyzes a sample leveraging on the information of the variants included in a `VCF` file, that is the standard format released by the 1000 Genomes Project [32] (`1KGP` from now on). To the best of our knowledge, `MALVA` is the first mapping-free tool able to call indels. Moreover, it proved to be the only such tool capable of handling the huge number of variants included in the latest version of the `VCF` released by the `1KGP`.

The rest of the paper is laid out as follows. In Section 2 we give some definitions and we describe the data structures used by `MALVA`. In Section 3 we thoroughly detail the methods. In Section 4 we show some implementation details and provide an experimental analysis of `MALVA` on real data to asses the tool’s performance against state-of-the-art tools. Finally, in Section 5, we draw conclusions and sketch possible future directions of our work.

## 2 Preliminaries

In this section we will describe the fundamental concepts that we will use in this document.

Let Σ be an ordered and finite alphabet of size *σ* and let *t* = *c*_1_,…,*c_k_*, where *c_j_* £ Σ for *j* = 1,…,*k*, be an ordered sequence of *k* characters drawn from Σ, we say that t is a *k*-mer. When a *k*-mer originates from a double stranded DNA, it is common to consider it and its reverse-complemented sequence as the same *k*-mer, and to say that the one that is lexicographically smaller among the two is the *canonical* one. In the following, we will abide by this definition and whenever we refer to a *k*-mer we implicitly refer to its canonical form. Moreover, to avoid *k*-mers being equal to their reverse-complement, we will only consider odd values of *k*.

A Bloom filter [1] is a probabilistic space-efficient data structure that represents a set of elements and allows approximate membership queries. The result of such queries may be a false positive but never a false negative. Bloom filters are usually represented as the union of a bitvector of length *m* and a set of *h* hash functions {H_1_,…, H_*h*_}, each one mapping one element of the universe to one integer in {1,…, *m*}. Using these data structures, the addition of an element *e* to the set is performed by setting to 1 the bitvector’s cells in positions {H_1_(*e*),…,H_*h*_(*e*)}, while testing if an element is in the set boils down to checking whether the same positions are all set to 1. Due to collisions of the hash functions, an element can be reported as present in the set even if it is absent. Nevertheless, the false positive rate of a Bloom filter of a set of *n* elements, with *h* hash functions and an array of *m* bits is 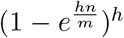; therefore, to increase the size of the Bloom filter decreases the false positive rate. Due to their simplicity and efficiency, Bloom filters have been applied to multiple problems in bioinformatics, such as representing de Bruijn graphs [2] and counting *k*-mers in a sample [23].

Let *B* be a bitvector, the rank_1_ function reports, for each position *i* ∈ {1,…, |*B*| + 1}, the number of 1s from the beginning of *B* to *i* (excluded); we refer to such value as rank_1_(*i, B*). Clearly, rank_1_(*i, B*) is not defined for *i* ≤ 0 and for *i* > |*B*| + 1, rank_1_(1, *B*) is 0, and rank_1_(|*B*| + 1, *B*) is the number of 1s in *B*. By using succinct support data structures and by a linear time preprocessing step, it is possible to answer rank_1_ queries in constant time for any position of the bitvector [36].

The difference between the genetic sequence of two unrelated individual of the same species is estimated to be smaller than 0.1% [35]; therefore, it is common to represent the DNA sequence of an individual as a set of differences from a *reference* genome. Indeed, thorough studies [3, 4, 6] of the variations across different individuals encode such information as a `VCF` (Variant Calling Format) file [8]. In the following, we will call *variant* the information encoded by a data line of a `VCF` file. Besides the genotype data, we are interested in the information carried by the second, fourth, fifth, and eighth field of a `VCF` line, namely: (i) field `POS` that is the position of the variant on the reference, (ii) field `REF` that is the reference allele starting in position `POS`, (iii) field `ALT` that is a list of alternate alleles that in some sample replace the reference allele, and (iv) field `INFO` that is a list of additional information describing the variant. From the latter list we will get the frequencies of reference and alternate alleles, which are needed to call the genotype of a given individual. We denote with `POS`(*v*), `REF`(*v*), `ALT`(*v*), `FREQ`(*v*), and `GTD`(*v*) the reference position, reference allele, list of alternate alleles, list of allele frequencies, and genotype data of a variant *v*, respectively.

The variants we take into account are SNPs (*i.e*., both `REF` and all the elements of `ALT` are single base nucleotides) and indels (`REF` and at least one element of `ALT` are not of the same length). Moreover, given an allele *a* (either reference or alternate) of some variant *v*, we refer to its sequence of nucleotides as `SEQ`(*a*), *i.e*., `SEQ`(*a*) is the string that represents *a*.

Let *R* be a reference genome and let `V` be a `VCF` file that describes all the known variants of *R*. Since the genotype data provides information on the alleles expressed in each genome, another way of thinking of a `VCF` file is as an encoding of a set of genomes 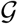. Each haplotype of the genomes in 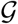 can be reconstructed by modifying *R* according to the genotype information associated to each variant. For ease of presentation, in the following we use the term genome and haplotype interchangeably, although each genome of a polyploid organism is composed of multiple haplotypes.

Let 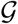 be the set of genomes encoded by a `VCF` file and let *a* be an allele of some variant *v*, we denote by 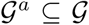 the subset of genomes that include *a*. We say that a variant *v* is *k-isolated* if there is no other known variant within a radius of ⌊*k*/2⌋ from the center of any of its alleles, as formally stated in the following definition.

### Definition 1

(*k*-isolated variant). *A variant v is k-isolated if, for all a* ∈ `ALLELES`(*v*) *and* 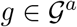, *there is no variant v*′ ≠ *v with an allele a*′ ∈ `ALLELES`(*v*′) *such that* 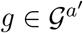 *and either* |`BEGIN`_*g*_ (*a*′) – `CENTER`_*g*_(*a*)| ≤ ⌊*k*/2⌋ *or* |`CENTER`_*g*_(*a*) – `END`_*g*_(*a*′)| ≤ ⌊*k*/2⌋, *where* `ALLELES`(*v*) = `REF`(*v*) ∪ `ALT`(*v*), `BEGIN`_*g*_(*a*) *is the position of the first base of a in g*, `END`_*g*_(*a*) *the position of the last base, and* `CENTER`_*g*_(*a*) *the position of the* 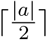-*th base of a in g*.

The procedure we will present in the next section is heavily based on the concept of *signature* of an allele. Intuitively, a signature of the allele *a* of a variant *v* is the *k*-mer centered in a in some genome *g* in 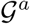. Note that, depending on the genomes encoded by the `VCF` file (specifically, if variants less than *k* bases apart are known), an allele might have multiple signatures. Moreover, if SEQ(a) is longer than *k* bases, the previous definition is not well formed, since there is no *k*-mer that can be centered in *a*. In this case, we define the signature of a as the set of its substrings of length *k*. The following definition formalizes the notion of signature of an allele.

### Definition 2

(Signature of an allele). *Let* 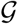 *be the set of all the genomes encoded by a* `VCF` *file* `V` *and let k be an odd positive value. Let v be a variant in* `V`, *let a be one of the alleles of v, and let* 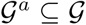 *be the set of the genomes that include a. If* `SEQ`(*a*) *is longer than k bases, we say that the signature of a is the set of all the substrings of length k of* `SEQ`(*a*). *If* `SEQ`(*a*) *is shorter than k bases, we say that* {*x*`SEQ`(*a*)*y*} *is the signature of a in a genome g in* 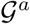 *if: (i) x*`SEQ`(*a*)*y is a k-mer, (ii)* 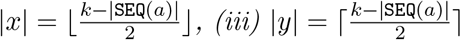, *(iv) x is a suffix of the sequence that precedes a in g, and (v) y is a prefix of the sequence that follows a in g*.

We will refer to the set of all the possible signatures of an allele *a* as `SIGN`(*a*) and we say that *k* is the *length of the signature*. An example of signatures of an allele is shown in Figure 1.

**Figure 1:**
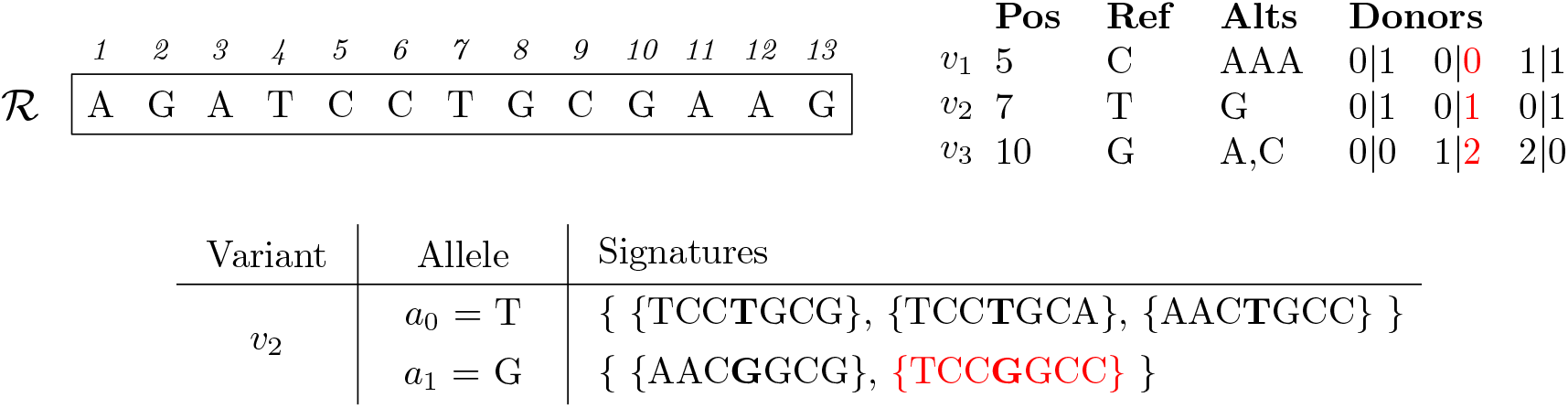
Signatures of the alleles of variant *v*_2_. 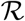 is the reference sequence and the table on the right is a `VCF` information associated to it, representing 3 variants: an indel (*v*_1_), a bi-allelic SNP (*v*_2_), and a multi-allelic SNP (*v*_3_). The last columns of the `VCF` file carry the genotype information of 3 individuals. The table at the bottom reports the signatures of each allele of variant *v*_2_. Note that there are only 5 signatures although 6 haplotypes are encoded by the `VCF` file since the second haplotype of the first and third individual are the same. We highlighted in red the genotype information associated to the second haplotype of the second genome and the corresponding signature.

In the following we will leverage on the definition of signature of an allele to detect its presence in an individual without mapping the reads to the reference genome. More precisely, we will analyze whether the *k*-mers of a given signature are present in the reads and use such information as an *hint* of the presence of the allele. Unlike other approaches [27], Definition 2 admits the presence of the alleles of multiple variants in a single signature, allowing `MALVA` to manage variants that are not *k*-isolated. Indeed, the set of signatures of an allele represents all the genomic regions where the allele appears in the genomes encoded by the `VCF` file.

## 3 Methods

In this section we will describe `MALVA`, the method we designed to genotype a set of known variants directly from a read sample. The general idea of `MALVA` is to use the frequencies of the signatures of a variant in the sample to call its genotype. The method works under the assumption that given a sample of reads from a genome with standard coverage depth, if an allele is included in the genome then at least one of its signatures must exist as substrings in multiple reads (depending on the coverage depth and the length of the signature). We leverage on this concept to genotype known variants directly from the input reads.

`MALVA` takes as input a reference genome, a `VCF` file representing all its known variants, and a read sample; it outputs a `VCF` file containing the most probable genotype for each variant. The main method is composed of four steps.

In the first step, `MALVA` computes the set of signatures of length *k_s_* of all the alternate alleles of all the variants in `VCF` and stores them in the set `ALTSIG`. In the same step, the signatures of the reference alleles are computed and stored in a second set named `REFSIG`. For each *k_s_*-mer *t* of a signature *s* two weights, one representing the number of occurrences of *t* in an alternate allele signature and one representing the number of occurrences of *t* in a reference allele signature, are stored. We will refer to these two values as 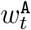 and 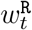, respectively.

We note that for small values of *k_s_* the probability that the *k_s_*-mers that constitute a signature appear in other regions of the genome is high. Since in the following steps `MALVA` exploits the signatures’ sets of the alleles of each variant to call the genotypes, the presence of conserved regions of the reference genome identical to some signature could lead the tool to erroneously genotype some variants. To get rid of a large amount of wrong calls, in the second step `MALVA` makes use of the context around the allele to distinguish its signatures from such regions. More precisely, if a *k_s_*-mer of a signature of an alternate allele appears somewhere in the reference genome, `MALVA` extracts the context of length *k_c_* (with *k_c_* > *k_s_*) covering the reference genome region and collects such *k_c_*-mers in a third set (`REPCTX`).

In the third step, `MALVA` extracts all the *k_c_*-mers from the sample along with the number of its occurrences. For each *k_c_*-mer *t_c_* that occurs *w* times in the sample, the *k_s_*-mer *t_s_* that constitutes the center of *t_c_* is extracted. If *t_s_* is found in `REFSIG`, 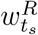 is increased by *w*. Moreover, if *t_c_* is not found in `REPCTX` and if *t_s_* is in `ALTSIG`, 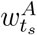 is increased by *w*. Otherwise, if *t_c_* is in `REPCTX`, 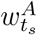 is not updated since, although its central *k_s_*-mer is identical to some *k_s_*-mer of a signature of an alternate allele of some variant, it is indistinguishable from another region of the genome not covering the variant. We note that when 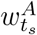 is not updated, our method might miss a variant in the donor and report a false negative, although for large values of *k_c_* this would rarely occur. The rationale behind this choice is to avoid biases due to *k_c_*-mers in conserved regions of the reference genome, preferring not to include an alternate allele in the output whenever ambiguities arise.

Finally, in the fourth step, `MALVA` uses the weights computed in the previous step to call the genotypes.

In the rest of this Section we will detail each one of the four steps of `MALVA`.

### Signature computation

The first step consists of building the signatures of the alleles of all the variants and adding them either to `ALTSIG`, if they are the signatures of an alternate allele, or to `REFSIG`, if they are the signature of the reference allele. If a variant *v* is *k_s_*-isolated, we build 1 + |`ALT`(*v*)| signatures, one for each allele of *v*. Otherwise, there are some genomes in 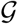 in which there is at least another allele of a variant that lays within a radius of ⌊*k_s_*/2⌋ nucleotides from the center of the allele of *v*. In practice, this means that we have to look at the genotype data of the variants within such radius: for each allele *a* of *v* we reconstruct the *k_s_* bases long portions of the genomes in 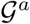 that constitute the signatures of *a*.

As pointed out in Definition 2, if |`SEQ`(*a*)| ≥ *k_s_*, the signature of *a* is the set of *k_s_*-mers that appear in `SEQ`(*a*). In this case we extract all such *k_s_*-mers and add them either to `REFSIG` or `ALTSIG`. Otherwise, if |*a*| < *k_s_*, we build the *k_s_* bases long substrings of each genome in 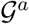 centered in *a* by scanning the `VCF` file and reconstructing the sequences according to the genotype information it includes. More precisely, let *a* be an allele of a variant *v* and let *V* = {*v*_1_,…,*v_n_*} be the set of variants such that, for all 1 ≤ *i* ≤ *n*: (i) *v_i_* ≠ *v*, (ii) there exists an allele *a_j_* in `ALLELES`(*v_i_*) such that *a* and *a_j_* are both included in some genome *g*, and (iii) either (`END`(*a_j_*) < `BEGIN`(*a*) and `CENTER`(*a*) – ⌊*k_s_*/2⌋ ≤ `END`(*a_j_*)) or (`END`(*a*) < `BEGIN`(*a_j_*) and `CENTER`(*a*) + ⌊k_s_/2⌋ ≥ `BEGIN`(*a_j_*)) in *g*.

Given *a*, we use the genotype information stored in the `VCF` file to retrieve the haplotypes in which it is included, *i.e*., a subset of the haplotypes in 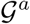, and build the set *V*. Using *V* we gather all the alleles that precede and succeed *a* in the selected haplotypes and we use them, together with the reference sequence, to reconstruct *on the fly* the *k_s_*-mer that covers *a*, by interposing reference substrings and allele sequences. Doing so, we don’t need to reconstruct the whole haplotypes but we only analyze and reconstruct the required *k_s_*-mers when needed.

Once all the *k_s_*-mers have been constructed, they are added to `REFSIG` if *a* is the reference allele, to `ALTSIG` if it is an alternate allele.

### Detection of repeated signatures

This step is aimed to detect and store in set `REPCTX` all the *k_c_*-mers of the reference sequence whose central *k_s_*-mer is included in some signature of some alternate allele, *k_c_* > *k_s_*. `REPCTX` will be used in a further step to discard alternate alleles that might be erroneously reported as expressed by `MALVA` only because they cannot be told apart from other identical regions of the reference sequence. To compute `REPCTX`, we extract all the *k_c_*-mers of the reference sequence and test whether their central *k_s_*-mer is in `ALTSIG`. If so, we add the *k_c_*-mer to `REPCTX` to report that the *k_s_*-mer is indistinguishable from some *k_s_*-mer that is included in the signature of an alternate allele. The set `REPCTX` is then used in the next step as illustrated below. An example comprising the first two steps in shown in Section B of the Appendix.

### Alleles’ signatures weights computation

In the third step, `MALVA` computes how many times the *k_s_*-mers of each signature appear in the dataset. First, `MALVA` extracts all the *k_c_*-mers of the read sample and tests their existence in `REPCTX` to check whether their central *k_s_*-mer cannot be told apart from some repetition in the reference genome. Then, given a *k_c_*-mer t_c_ that occurs *w* times in the read sample, the *k_s_*-mer *t_s_* that constitutes its center is extracted. If *t_s_* is found in `REFSIG`, *i.e., t_s_* is the signature of the reference allele of some variant, the weight 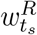 is increased by *w*. Moreover, if *t_c_* is not found in `REPCTX` and *t_s_* is in `ALTSIG`, *i.e*., *k_s_*-mer *t_s_* is uniquely associated to an alternate allele of some variant, the weight 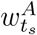 is increased by *w*. Conversely, if *t_c_* is in `REPCTX`, 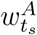 is not updated. The last scenario happens when *t_s_* is identical to the signature of an alternate allele of some variant (indeed, *t_s_* is in `ALTSIG`), but even the enlarged context *t_c_* (and consequently *t_s_*) appears somewhere else in the reference genome.

### Genotype calling

In the last step, `MALVA` uses the allele frequencies stored in the `INFO` field of the `VCF` file and the weights of the signatures computed in the previous step to call the genotype of each variant. To this aim, we extend the approaches proposed in the literature for bi-allelic variants [30, 34] to multi-allelic variants.

Let *v* be a variant with *n* − 1 alternate alleles. The number of possible distinct genotypes is 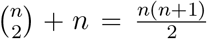, that is one *homozygous reference* genotype, 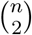 *heterozygous* genotypes, and *n* − 1 *homozygous alternate* genotypes. We will refer to the homozygous reference genotype as *G*_0,0_, to the heterozygous genotypes as *G_i,j_* with 0 ≤ *i* < *j* ≤ *n* − 1, and to the homozygous alternate genotypes as *G_i,i_* with 1 ≤ *i* ≤ *n* − 1. Following well-established techniques [30, 22, 18], we compute the likelihood of each genotype *G_i,j_* by means of the Bayes’ theorem. Given the observed coverage *C*, we compute the posterior probability of each genotype as:

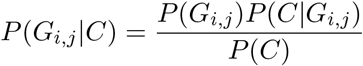

that, by the law of total probability, can be expressed as:

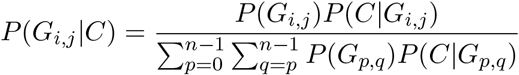

To calculate this probability, we compute the *a priori* probabilities of each genotype *G_i,j_*(*P*(*G_i,j_*)) and the *conditional probability* of the observed coverage given the considered genotype (*P*(*C|G_i,j_*)). The Hardy-Weinberg equilibrium equation ensures that for each variant *v*, (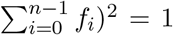, where *f_i_* = `FREQ`(*v*)[i], *i.e*., the frequency of the *i*-th allele of *v*. We recall that `FREQ`(*v*) is stored in the `INFO` field of the `VCF` file. The *a priori* probability of each genotype *G_i,j_* is therefore computed as follows:

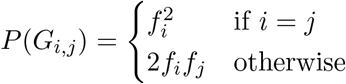

To compute the conditional probability *P*(*C|G_i,j_*), it is first necessary to compute the *coverages* of the alleles of the variant. Without loss of generality, let *a*_0_ be the first allele of the variant, *i.e., a*_0_ is the reference allele with index 0. We recall that `SIGN`(*a*_0_) is the set of signatures of allele *a*_0_ and that each signature is a set of one or more *k*-mers. We also recall that, in the previous step, for each *k*-mer *t* that belongs to some signature we computed two weights, namely 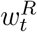 and 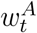. Given a signature *s* ∈ `SIGN`(*a*_0_), we define its weight as the mean of the weights associated to the *k*-mers it contains, *i.e*., 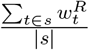 where |*s*| denotes the number of *k*-mers contained in signature *s*. Since the same allele may exhibit more signatures, we define the coverage *c*_0_ of allele *a*_0_ as the maximum value among the weights of its signatures, *i.e*., 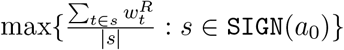. This formula can be easily modified to compute the coverage of an alternate allele (*c_i_* for *i* ≥ 1) by switching 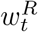 with 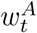. The coverage *c_i_* of an allele *a_i_* of a variant is thus computed as follows:

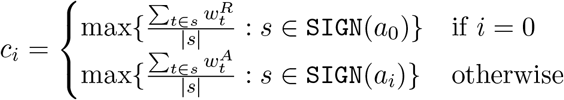

By extending the approach adopted in [30], we consider each *P*(*C|G_i,j_*) to be multinomially distributed. Given a homozygous genotype *G_i,i_*, we assume to observe the *i*-th allele, which is the correct one, with probability 1 − *ε* (where *ε* is the expected error rate) whereas the other *n* − 1 alleles (the erroneous ones) with probability 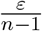 each. Hence, we compute the conditional probability of an homozygous genotype as:

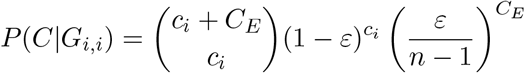

where *C_E_* is the total sum of the coverages of the erroneous alleles, *i.e*., 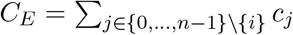. For what concerns heterozygous genotypes, we assume to observe the correct alleles, *i.e*., the *i*-th and the *j*-th allele, with equal probability 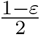 whereas the other *n* − 2 erroneous alleles with probability 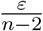 each. We compute the conditional probability of an heterozygous genotype as follows:

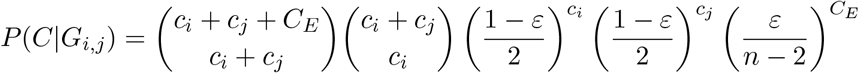

where, again, *C_E_* is the sum of the coverages of the erroneous alleles, *i.e*., 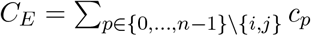.

Finally, after computing the posterior probability of each genotype, `MALVA` outputs the genotype with the highest likelihood.

## 4 Implementation and Experiments

In this section we will describe the implementation details of `MALVA` and we will provide an experimental analysis on real data. All the analyses were performed on a 64 bit Linux (Kernel 4.4.0) system equipped with four 8-core Intel ^®^ Xeon 2.30GHz processors and 256GB of RAM.

### Implementation details

`MALVA` is implemented in `C++` and it is freely available at https://github.com/AlgoLab/MALVA. Bloom filters were implemented as the union of a bitvector and a single hash function `H`. Although it is not conventional, in most cases to use a single hash function has similar results as using multiple ones, as noticed by other authors [33, 34]. To check this claim, while developing the tool we tested whether using multiple hash functions would improve the results by extending the Bloom filters to count-min sketches [7]. As expected, the deterioration of the performance far outweighted the gain in precision and recall (that was less than 0.1%). Moreover, to use a single hash function allows us to store 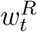 and 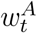 efficiently for each *k*-mer *t*. Indeed, note that once all the signatures of all the alternate alleles have been added to `ALTSIG`, the latter is only used to check whether some *k_s_*-mer is part of a signature, *i.e*., it becomes static. By representing `ALTSIG` as a Bloom filter *B*_ALTSIG_ we can create an integer array `CNTS` of size rank_1_(|*B*_ALTSIG_| + 1,*B*_ALTSIG_) to store the weights of each *k*-mer compactly and, if a *k*-mer *t* of a signature *s* is in `ALTSIG` (*i.e*., if *B*_ALTSIG_[`H`(*s*)] = 1) we can access its weight by accessing `CNTS`[rank_1_(`H`(*s*), *B*_ALTSIG_)]. In a nutshell, after adding all the alternate alleles to *B*_ALTSIG_, we *freeze* it, build a rank data structure over it, compute the number of ones, and create the `CNTS` array of the correct size. Similarly, we implemented `REPCTX` as a Bloom filter *B*_REPCTX_ using a single hash function. Conversely, `REFSIG` was implemented as a simple hash table, because the number of elements it stores is usually smaller than the number of elements stored in `ALTSIG`. The bitvectors, the rank data structure, and the `CNTS` array were implemented using the `sdsl-lite` library [12]. We pose an upper limit of 255 to the value of each cell of the `CNTS` array, so as to store each counter using only 8 bits.

Finally, instead of scanning all the *k_c_*-mers in the read sample, we used `KMC3` [10] to efficiently extract them and counting their occurrences. Therefore, in step 3 `MALVA` parses the output of `KMC3` and updates the counts for each *k_s_*-mer accordingly.

### Experiments

We performed an experimental analysis on real data to evaluate the real feasibility of our method, comparing `MALVA` to one mapping-free method and to two different alignment-based pipelines. Among the mapping-free methods proposed in literature we chose `VarGeno`, as it is an improved version of `LAVA` that provides better efficiency and accuracy [34]. For completeness, we included in our evaluation the two most widely used alignment-based pipelines, denoted by `bcftools` and `GATK`, respectively. The former consists of an alignment step performed with `BWA-MEM` [19] followed by a variant discovering step performed using `bcftools` [18]. The latter consists of an alignment step performed with `BWA-MEM` and a variant discovering step performed with `GATK` [22], as recommended by the *GATK Best Practices* [9].

`MALVA` was run setting *k_s_* equal to 47, *k_c_* equal to 53, *ε* equal to 0.1%, Bloom filters size equal to 8*GB*, and considering the a priori frequencies of the alleles of the `EUR` population, since the individual under analysis is part of it.

We tested the tools using the Illumina WGS dataset of the well-studied NA12878 individual provided by the Genome In A Bottle (GIAB) consortium [38]. We chose this individual because the variant calls provided are highly reliable and can be effectively used to assess the precision and the recall of the considered methods. We downloaded the alignments of its 30x downsampled version and used `SAMtools` [20] to extract the corresponding `FASTQ` file, obtaining 696,168,435 150bp-long reads. As reference genome and set of known variants, we used the GRCh37 primary assembly and the `VCF` files provided by Phase 3 of the `1KGP` [32]. These `VCF` files contain a total of 84,739,838 variants, the phased genotype information of 2504 individuals, and the a priori frequency of each allele of each variant of 5 populations. We note that `VarGeno` requires a different formatting of the fields describing the a priori frequencies of the alleles than the ones in the `VCF` file provided by the `1KGP`. Thus, we formatted the input files as required before running `VarGeno`.

`VarGeno` could not complete the analysis of this dataset, from now on denoted by `FullGenome`, on our server. To test whether `VarGeno` crashed due to excessive memory usage, we tried to run it on the same instance on a cluster with 1TB of RAM, but nevertheless it could not complete the analysis, crashing after 20 minutes. In order to include `VarGeno` in our evaluation, we chose 12 chromosomes to create a smaller dataset, denoted by `HalfGenome`, that thus contains some half of the variants and the reads of the `FullGenome` dataset.

Each tool was evaluated in terms of variant calling accuracy and efficiency (wall time and memory usage). We note that some steps of the previously cited tools can use multiple threads to improve the time performance (namely, `KMC3` for `MALVA`, `BWA`-MEM for `bcftools` and `GATK`, and the variant discovery steps of `GATK`). Whenever we had this choice, we provided 4 threads to each tool. We used `hap.py` [15], the tool developed for the evaluation of variant callers in the recent *PrecisionFDA Truth Challenge*^1^, and the /usr/bin/time system tool to gather the required data.

Table 1 shows the results obtained by the considered tools on both the `FullGenome` and the `HalfGenome` datasets. We point out that `hap.py` computes precision and recall considering only non-reference `VCF` records (*i.e*., non 0/0 calls).

**Table 1:** Accuracy and efficiency results on the `HalfGenome` and `FullGenome` datasets. For each dataset, we reported the values of Precision (P) and Recall (R) obtained by the considered tools on both SNPs and indels. The efficiency results are shown in terms of wall clock time and peak memory usage. `VarGeno` could not complete the analysis of the `FullGenome` dataset, thus we did not report its results on this dataset.

| Dataset | Tool | P<sub>SNP</sub> | R<sub>SNP</sub> | P<sub>INDEL</sub> | R<sub>INDEL</sub> | Time (hh:mm) | RAM (GB) |
| --- | --- | --- | --- | --- | --- | --- | --- |
| HalfGenome | MALVA | 92.9% | 90.7% | 85.3% | 80.9% | 03:19 | 30 |
|  | VarGeno | 97.5% | 88.1% | 39.5% | 0.1% | 02:31 | 52 |
|  | bcftools | 91.2% | 94.8% | 44.9% | 55.4% | 24:35 | 6 |
|  | GATK | 91.7% | 95.1% | 53.2% | 79.9% | 34:43 | 32 |
| FullGenome | MALVA | 91.6% | 89.7% | 84.4% | 80.1% | 07:31 | 37 |
|  | VarGeno | - | - | - | - | - | - |
|  | bcftools | 91.6% | 94.4% | 44.7% | 54.6% | 54:23 | 9 |
|  | GATK | 92.0% | 94.7% | 53.2% | 79.2% | 73:36 | 33 |

As expected, `MALVA` and `VarGeno` are one order of magnitude faster than `bcftools` and `GATK`. Indeed, `MALVA` and `VarGeno` required 3.5 and 2.5 hours to analyze the `HalfGenome` dataset, respectively, while `bcftools` and `GATK` required 24.5 and 34.5 hours. We note that half of the time required by `bcftools` and one third of the time required by `GATK` was spent running `BWA-MEM`, that completed its task in 12.5 hours (using 4 threads). The same trend applies to the analysis of the `FullGenome` dataset, on which each tool required roughly twice the time required on the `HalfGenome` dataset. A qualitative representation of each tool’s running times is shown in Figure 2.

**Figure 2:**
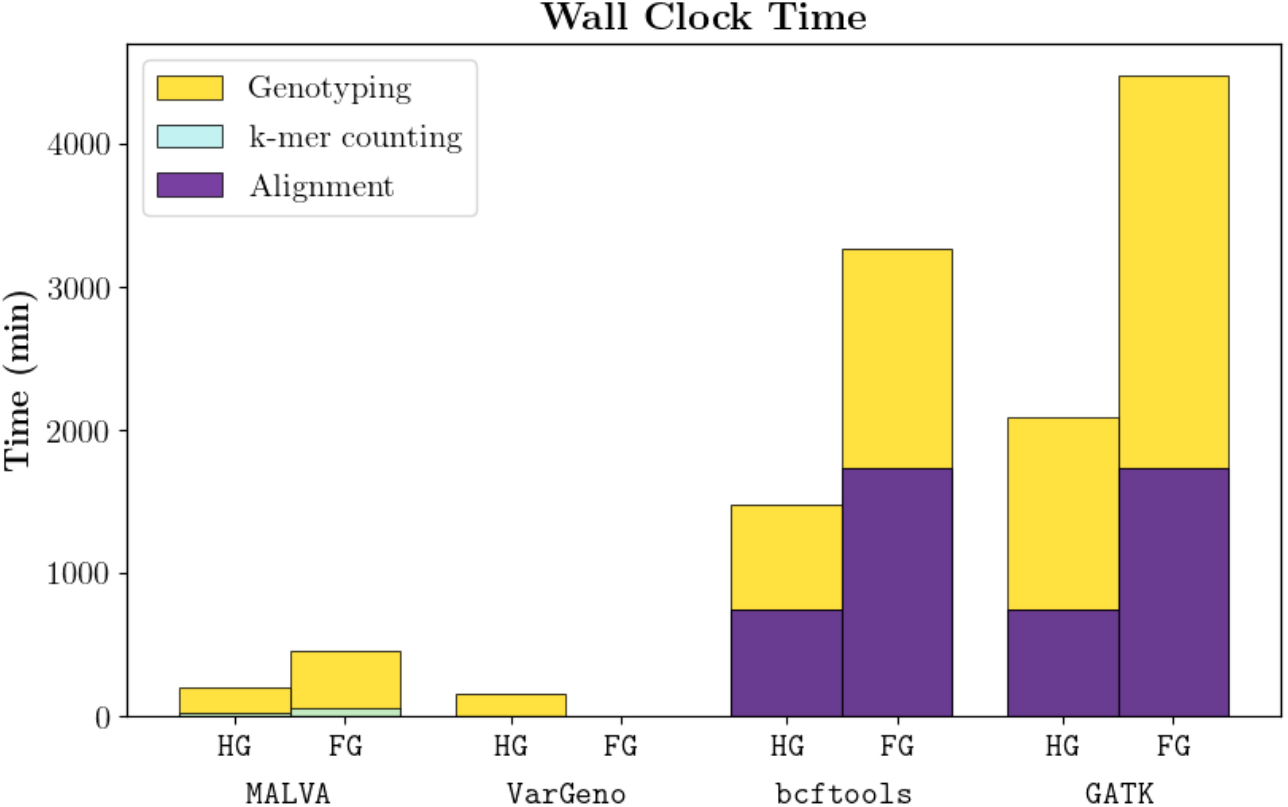
Times required by each tool to analyze both datasets, partitioned by steps performed. For ease of presentation, we denoted the `FullGenome` dataset as `FG` whereas the `HalfGenome` dataset as `HG.` Note that we did not include `VarGeno` running time on the `FullGenome` dataset since it crashed after 20 minutes.

For what concerns the memory usage, `bcftools` proved to be the least memory intensive approach, requiring less than 10GB of RAM on both datasets to map the reads and less than 1GB of RAM to call the variants. `MALVA` and `GATK` showed similar memory requirements, with `GATK` showing almost no difference between the two analyses and `MALVA` increasing the memory consumption by only 23% for the bigger dataset. `VarGeno` required slightly less than twice the amount of memory required by `MALVA` on the `HalfGenome` dataset.

Precision and recall of all the tools varied little over the two datasets, proving that the number of variants and reads only slightly affects their accuracy. As expected, `bcftools` and `GATK` achieved the best recall for non-homozygous reference SNPs due to the mapping step, that provides a more precise coverage of the alleles and allows to better discern repeated regions of the reference genome. `VarGeno` achieved the lowest recall obtaining 2% less recall than `MALVA`, that in turn called correctly 90.7% of the SNPs. On the other hand, `MALVA, bcftools`, and `GATK` achieved comparable precision on SNPs, whereas `VarGeno` obtained the highest one. Overall, on non-homozygous reference SNPs, `VarGeno` seems to be the most conservative tool among those tested, as it prefers not to call SNPs when there is any uncertainty. On the contrary, `MALVA`, in avoiding the loss of any potentially interesting information, deliberately prefers to detect any potential alternate allele in the donor, at the cost of a slight loss in precision

Remarkably, on indels `MALVA` obtained significantly better recall than `bcftools` and better precision than any other tool. As expected, since the method of `VarGeno` is not designed to manage indels, it was only able to genotype a negligible percentage of them. `bcftools` showed a very low precision and recall on indels, whereas `GATK` achieved a recall similar to `MALVA` but a low precision. The low precision achieved by the alignment-based tools is mainly due to the difficulties in aligning reads that overlap with indels.

Overall, `MALVA` proved to be an accurate and efficient alternative to mapping-based pipelines for variant calling, achieving good results both on SNPs and indels. The experimental evaluation shows the usefulness of the formalization of signature of an allele, of the extension to multi-allelic SNPs and indels, and of the ability to manage variants in dense genomic regions. A more in-depth comparison of `MALVA` and `VarGeno` is provided in Section A of the Appendix.

## 5 Conclusions and Future Directions

In this article, we presented `MALVA`, the first efficient mapping-free genotyping tool that is able to handle multi-allelic variants and indels. We compared `MALVA` with `VarGeno`, the state-of-the-art mapping-free genotyping tool, showing that our method is less memory intensive, achieves better recall, handles dozens of millions of variants effectively, and provides correct genotypes even for indels. We also compared our tool with two variant discovery pipelines, namely `GATK` and `bcftools`, showing that `MALVA` is an order of magnitude faster while achieving better accuracy on indels and similar accuracy on SNPs.

`MALVA` proved to be able to efficiently manage a huge amount of variants like those provided by the 1000 Genome Project (80 millions of variants) and to handle multi-allelic variants and indels. These fundamental features allow our method to exploit the whole information in input, without filtering out any data that might be crucial in successive analyses. Most notably, `MALVA`’s ability to genotype indels allows to apply mapping-free techniques to many clinical contexts, including screens for genetic predispositions for disease linked to the presence of indels [28, 37].

Future steps will be devoted to improving the efficiency of `MALVA` by exploiting the parallel architecture of modern machines, and to extending the method to genotype trios.

## A Comparison of MALVA and VarGeno output

To assess whether the tools under analysis produce some systematic error, we considered the `HalfGenome` dataset and we performed a more in-depth analysis of the SNPs in the `VCF` output by `MALVA` and `VarGeno`. For each tool, Figure 3 reports the number of correct genotypes output, grouping them in *homozygous reference* (*i.e*., 0|0), *heterozygous reference* (*i.e*., 0|1, 0|2, and so on.), *homozygous alternate* (*i.e*., 1|1, 2|2, and so on), and *heterozygous alternate* (*i.e*., 1|2, 1|3, 2|3, and so on). As stated in Section 4, we recall that the precision and recall output by `hap.py` do not consider homozygous reference genotypes, thus the analysis we present in this section allows us to better understand the behavior of the tools. Since `VarGeno` is not able to manage indels, we decided not to include them in this analysis.

Consistently with the precision and recall results of `hap.py, MALVA` detects between 5% and 10% more correct variants than `VarGeno` in all classes, at the cost of producing more erroneous calls. We note that overall `VarGeno` filters out 2,004,259 of the 39,796,878 SNPs in the truth.

Both tool show similar pattern in erroneous calls. More precisely erroneously genotyped homozygous reference variants were mostly genotyped as heterozygous reference and, vice-versa, erroneously genotyped heterozygous reference variants were mostly genotyped as homozygous reference. On the other hand, erroneous homozygous alternate variants in the donor were mostly genotyped as heterozygous reference by `VarGeno` whereas `MALVA` evenly distributed the errors between homozygous reference and heterozygous reference calls. Finally, erroneous heterozygous alternate variants in the donor were mostly genotyped as homozygous alternate variants by `MALVA`, meaning that the method proposed in this paper was able to detect the fact that the allele was not the reference allele but it called one of the two alternate alleles of the donor erroneously. Figure 3 shows a comparison between real genotype (provided by the 1000 Genomes Project) and genotype called by `MALVA` and `VarGeno` on SNPs. Figure 4 shows the same data row-normalized.

**Figure 3:**
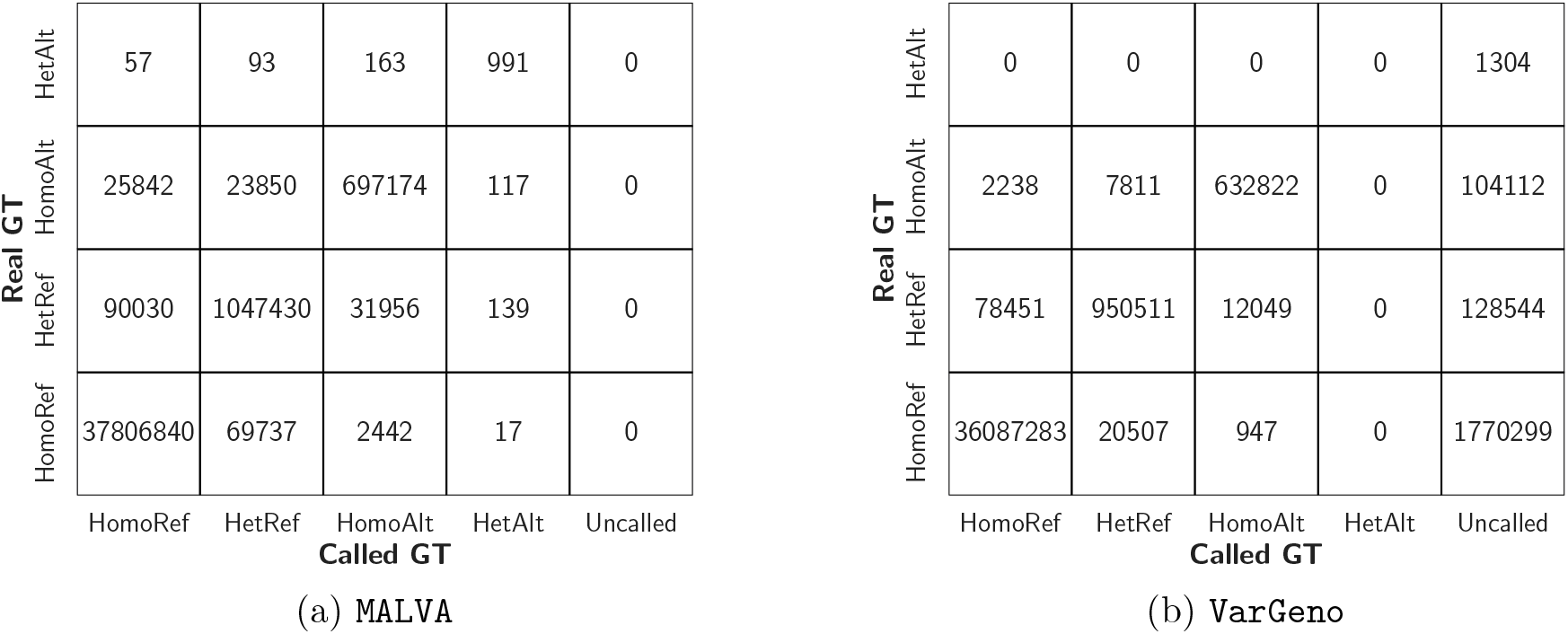
Comparison between real genotype (provided by the 1000 Genomes Project) and genotype called by each tool. HomoRef stands for Homozygous Reference, HetRef stands for Heterozygous Reference, HomoAlt stands for Homozygous Alternate, HetAlt stands for Heterozygous Alternate, and Uncalled means that the given variant was not called by the tool.

**Figure 4:**
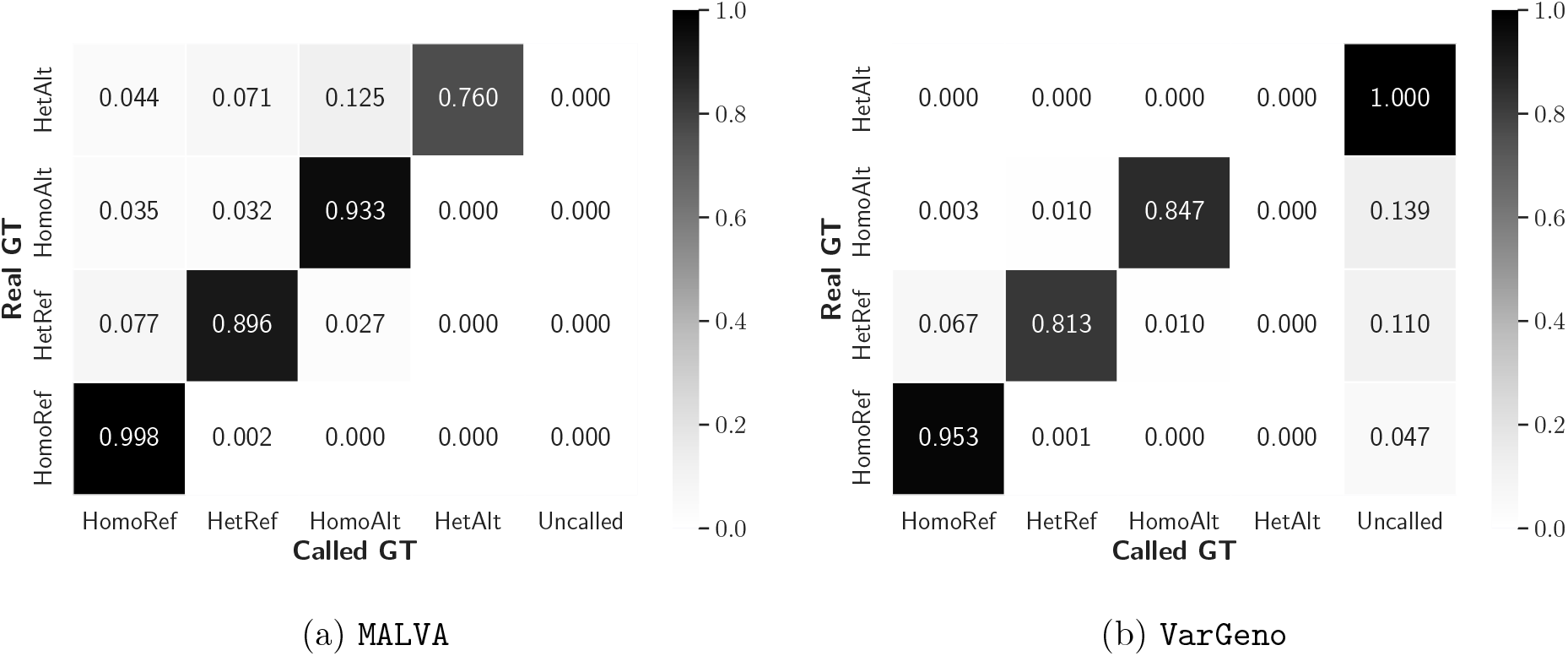
Comparison between real genotype (provided by the 1000 Genomes Project) and genotype called by each tool, normalized by rows.

Overall, the errors produced by both tools were “partial” errors in the sense that they rarely mis-call both alleles of the donor.

## B Example *k*-mers weight computation

In this section we present an example of computation of the weights associated with the signatures’ *k_s_*-mers. Figure 5 shows an example composed of three variants and two reads. In this example the values of *k_s_* and *k_c_* are set to 7 and 11, respectively. Subfigure (*a*) shows the 26-bases long reference sequence. Subfigure (*b*) reports on the left two bi-allelic variants (*v*_1_ and *v*_2_) and one multi-allelic variant (*v*_3_), and on the right the signatures of each allele of *v*_2_. Subfigure (*c*) shows the elements of `ALTSIG` and `REFSIG` related to *v*_2_. We note that the second signature in `ALTSIG` is composed of a single *k_s_*-mer (*t_s_*, equal to `TCCGGCG`) that appears in the reference genome, starting from position 17. Thus, the *k_c_*-mer starting in position 15 and ending in position 25 (*t_c_*, equal to GATCCGGCGAA) is added to `REPCTX`. Subfigure (*d*) shows two 11-bases long reads including *t_s_*, extracted from position 3 and 15 of the donor. Clearly, only *r*_1_ should contribute to the detection of the alternate allele of *v*_2_ in the donor, since *r*_2_ was sequenced from another position of the genome (*i.e*., 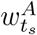 should be equal to 1 in this case). To this aim, `REPCTX` comes to an aid; indeed, when analyzing *r*_1_ the *k_c_*-mer covering *t_s_* is extracted (*i.e*., the whole read) and its inclusion in `REPCTX` is tested. Since TATCCGGCGTA is not in `REPCTX` and *t_s_* is in `ALTSIG`, 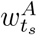 is increased by one. On the other hand, since GATCCGGCGAA is in `REPCTX`, the occurrence of *t_s_* in *r*_2_ is not considered in 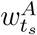, thus avoiding to erroneously overestimate the frequency of allele *a*_1_ of *v*_2_.

We note that on one hand this approach allows us to avoid overestimating the frequencies of some alternate allele but, on the other hand, it produces two major side effects. The first one is that some allele might be underestimated by `MALVA`; indeed, if the *k_c_*-mer covering an alternate allele in a donor is equal to a *k_c_*-mer in the genome it will not be detected. The second side effect is that `MALVA` might overestimate the frequency of some allele due to identical signature. Indeed, suppose that the signature of some alternate allele *a_i_* of another variant *v_j_* ≠ *v*_2_ is equal to the signature of alternate allele *a*_1_ of variant *v*_2_. It is obvious that the weights of the *k_s_*-mers of the two signatures will be identical and that the occurrences of both the alleles will concur towards their final value, overestimating it.

Although the two side effects pose some limit to the method proposed in this paper, they arise rarely and we think they are a fair price to pay to avoid biases introduced by the reference genome.

**Figure 5:**
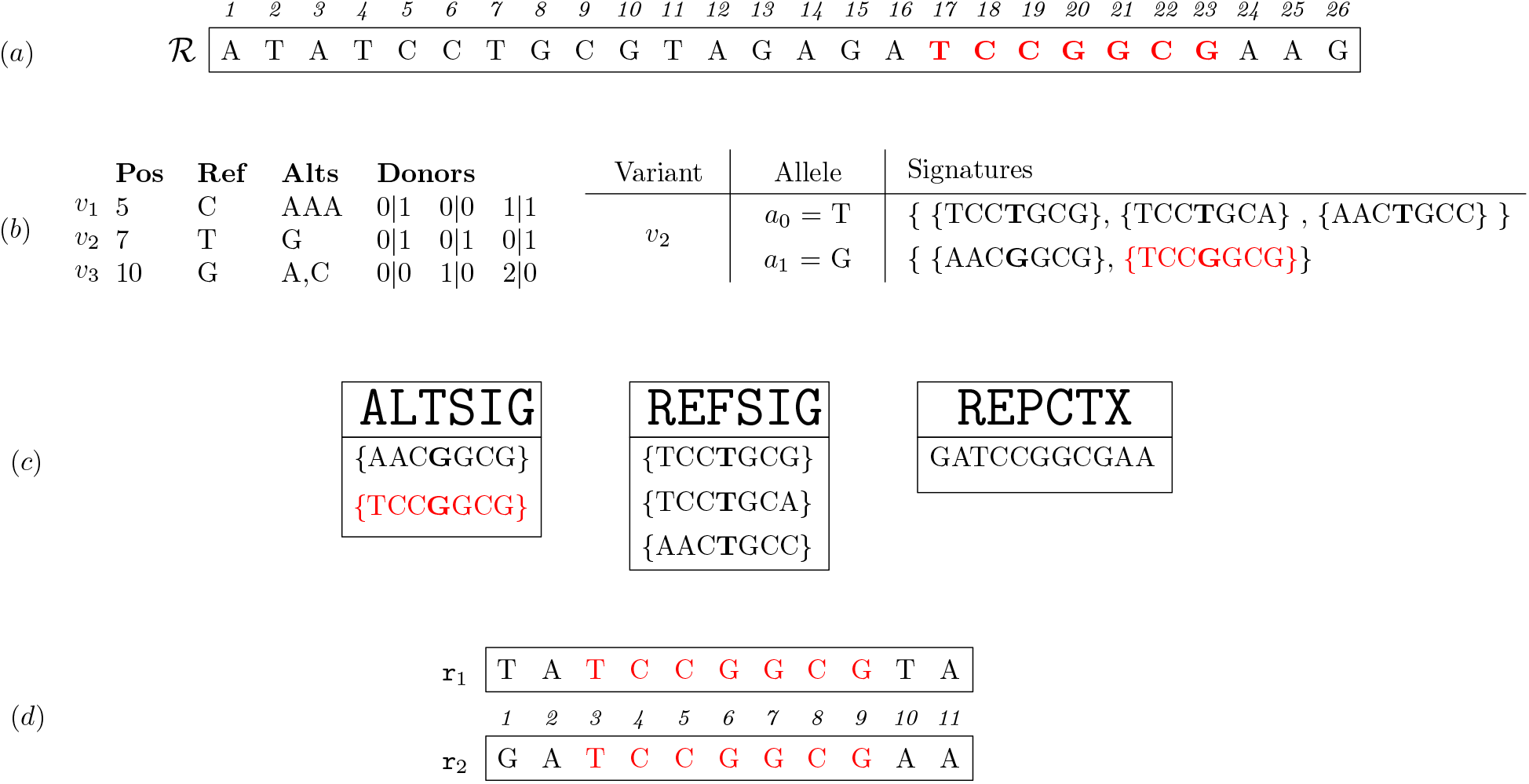
Example with 3 variants and two reads. Subfigure (*a*) shows a reference genome of 26 bases, Subfigure (*b*) reports 3 variants and the signatures of each allele of variant *v*_2_, Subfigure (*c*) reports the subsets of `ALTSIG`, `REFSIG`, and `REPCTX` including the elements related to *v*_2_, and Subfigure (*d*) presents two reads of length 11.

## Footnotes

1 https://precision.fda.gov/challenges/truth

